# Surface tension determines tissue shape and growth kinetics

**DOI:** 10.1101/456228

**Authors:** S. Ehrig, C. M. Bidan, A. West, C. Jacobi, K. Lam, P. Kollmannsberger, A. Petersen, P. Tomancak, K. Kommareddy, F. D. Fischer, P. Fratzl, John W. C. Dunlop

## Abstract

The growth of tissue is an essential process controlling morphogenesis and regeneration of organs [1]. In general tissue forming cells are interactive and motile [2], which can give rise to emergent physical properties such as viscous fluid behaviour as has been shown for epithelial monolayers during embryogenesis [3, 4], and for cell-agglomerates [5] with a measurable surface tension [6]. However, the mechanical integrity of tissues is provided by extracellular matrices (ECM) that turn tissues into solids with well-defined elastic properties [7]. Paradoxically, it has been shown by in-vitro experiments that even osteoid-like tissue with large amounts of ECM grows according to rules reminiscent of fluid behavior [8, 9]. Motivated by this conundrum, we show here quantitatively, by constraining growing tissues to surfaces of controlled mean curvature, that osteoid-like tissues, develop shapes similar to the equilibrium shapes of fluids [10]. In particular, for geometries with rotational symmetry, the tissue stays bounded by Delaunay surfaces and grows with rates depending on surface curvature. Actin stress-fibre patterns at the tissue surface suggests that cell contractility is responsible for generating the necessary surface stresses. This indicates that continuous remodeling of the solid matrix combined with the contractility of bone forming cells provide sufficient effective fluidity and surface stress required for a fluid-like behavior of the growing tissue at the time scale of days to weeks. Our work demonstrates that morphogenesis shares fundamental physical principles with fluid droplets as first suggested in D’Arcy Thompson’s seminal work ‘On Growth and Form’ [11].

Single cells respond to the dimensionality of their environment [12, 13], with notable differences between cells on flat surfaces and those surrounded by a 3D environment [14]. In vivo, cells are unlikely being constrained to flat surfaces [15] but, as long as the curvature is small, cells may behave as if the surface is flat [16]. For cell agglomerates and tissues the situation is different as cells can then mechanically interact with each other as well as with their physical environment [17-19]. They may even collectively sense and respond to macroscopic surface curvature [8, 9]. Indeed, the geometry of the environment strongly influences cell-behavior, since it determines the spatial distribution of force patterns that cells sense and transmit. Previous studies on the role of surface curvature on cell and tissue behavior focused on surfaces where one of the principal curvatures is zero [8, 20, 21] or was not quantitative due to the complexity of the scaffolds [22]. Here we address this problem by growing tissue on scaffolds with rotational symmetry and constant mean curvature. These scaffolds were obtained by shaping liquid polydimethylsiloxane (PDMS) through surface tension using a method adapted from Wang and McCarthy [23]. By constraining a polymer drop between two solid disks a capillary bridge (CB) with constant mean curvature (Supplementary Material Fig S4) forms. Its mean curvature can be precisely tuned by varying bridge height, liquid volume and disk diameters (Supplementary Material Fig S4). If tissue growing on such a surface has fluid-like behavior, it must morph into shapes delimited by “Delaunay surfaces” [10].

In order to investigate if the growing tissue behaves like a liquid, osteoid-like tissue was grown on these PDMS scaffolds using a pre-osteoblastic cell-line (MC3T3-E1) which synthesizes a collagen-rich extracellular matrix. Tissue growing on the curved surface was pinned to the flat edges of the bridge (Fig. 1A; red circle). In addition, the tissue growing in between the CB and the sample holder displayed a moving contact line. Both observations are reminiscent of liquid wetting. Interestingly, for a constant height of the CBs, the average thickness of tissue formed after about one month of culture was observed to decrease with increasing neck radius of the CB (Fig. 1 B, C, F-I and J-L). Additionally the tissue surfaces were shown to be rotational symmetric using 3D Light Sheet Florescence Microscopy (LSFM) imaging of fixed tissues (Fig. 1B-E, Suppl. film). This rotational symmetry allowed us to estimate tissue volume and mean curvature from 2D images. To explore the behavior of the growing tissue, we plotted mean curvatures and surface areas of the tissue as a function of total volume (CB volume plus tissue) and compared this to theoretical predictions for a liquid drop adhering to the scaffold using Brakke’s Surface Evolver [24](Fig. 2A, B). Liquid-like behavior was demonstrated also for CBs with smaller end-spacings (Fig. 2C-D) that cover a larger range of mean curvature. Remarkably, our experimental results perfectly match the theoretical predictions of the Laplace-Young law (see suppl. Material) providing strong experimental evidence that growing osteoid-like tissues behave like fluids at time scales relevant for growth.

**Figure 1:**
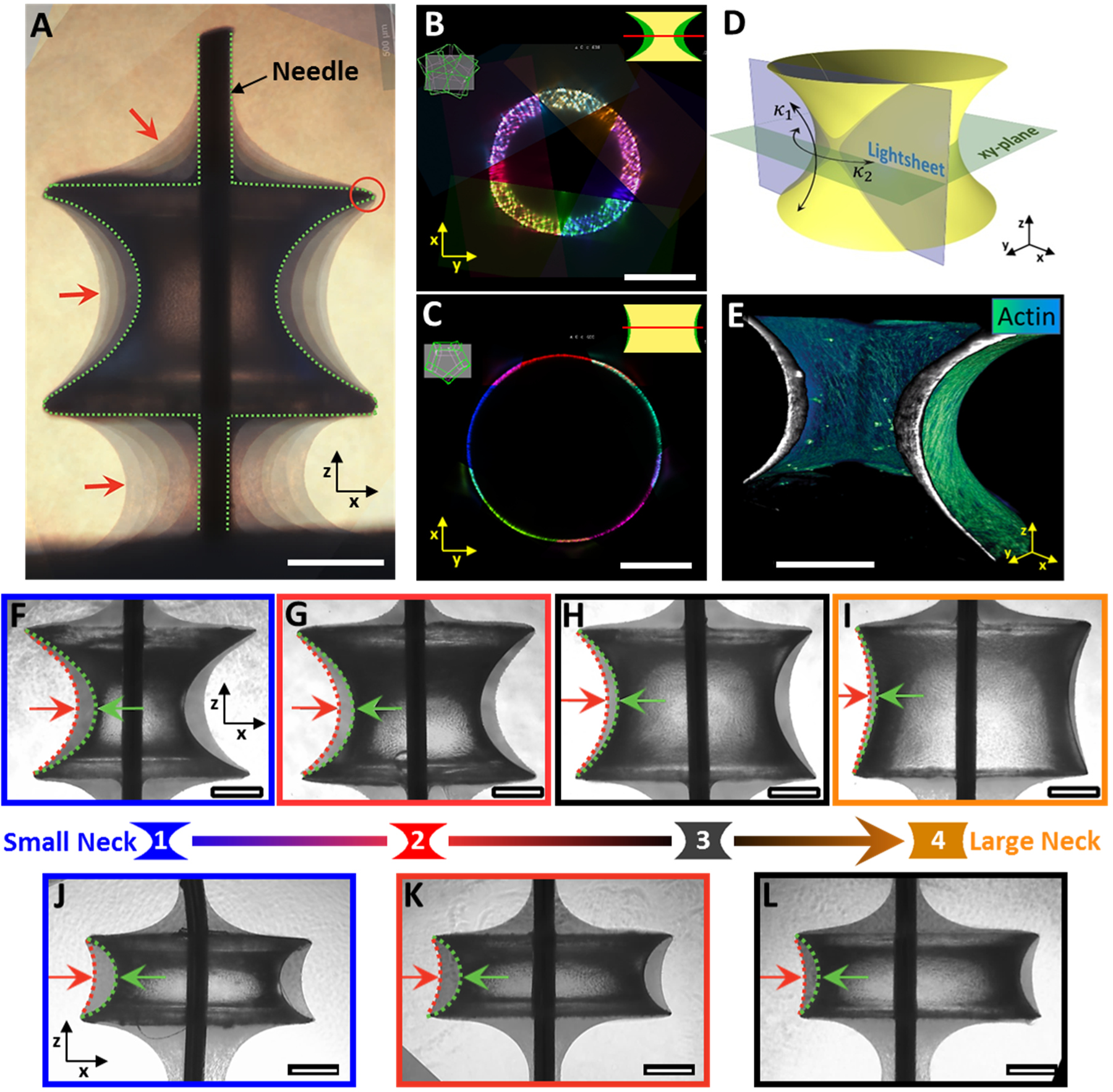
Capillary bridge geometry as a means to control 3D tissue growth. (A) Composition of phase-contrast images of tissues grown on a capillary bridge (CB) taken after 4, 7, 21, 32, 39, and 47 days. The tissue is pinned at the edges of the capillary bridge (red circle), and shows a moving contact line in-between scaffold and Teflon holder, reminiscent of a liquid. Dashed line indicates capillary bridge surface and red arrows point towards the tissue-medium interface. Scale bar: 500µm. (B) and (C) are radial slices at the neck for two different capillary-bridge sizes of 1.1µ (B) and 2.8µl (C) initial volume obtained with Light-Sheet Fluorescence Microscopy (LSFM) from 5 different views. (D) Sample geometry and orientation of the light-sheet. *κ*_1_ and *κ*_2_ are minimum and maximum principal curvatures. (E) 3D rendering of actin fibres on the sample shown in (F) color-coded according to fluorescence intensity. (F-I) phase contrast images of tissues grown on four different capillary bridge surfaces with initial volumes of 1.1μl (F), 1.6μl (G), 2.2μl (H), and 2.8μl (I). (J-L) capillary bridge surfaces with initial volumes of 1.1 μl (J), 1.3μl (K), and 1.5μl (L). Sample neck-size increases from left to right. Green arrow indicates interface of initial shape, and red arrow indicates position of tissue-medium interface after 32 days. Scale bars are 400μm.

**Figure 2:**
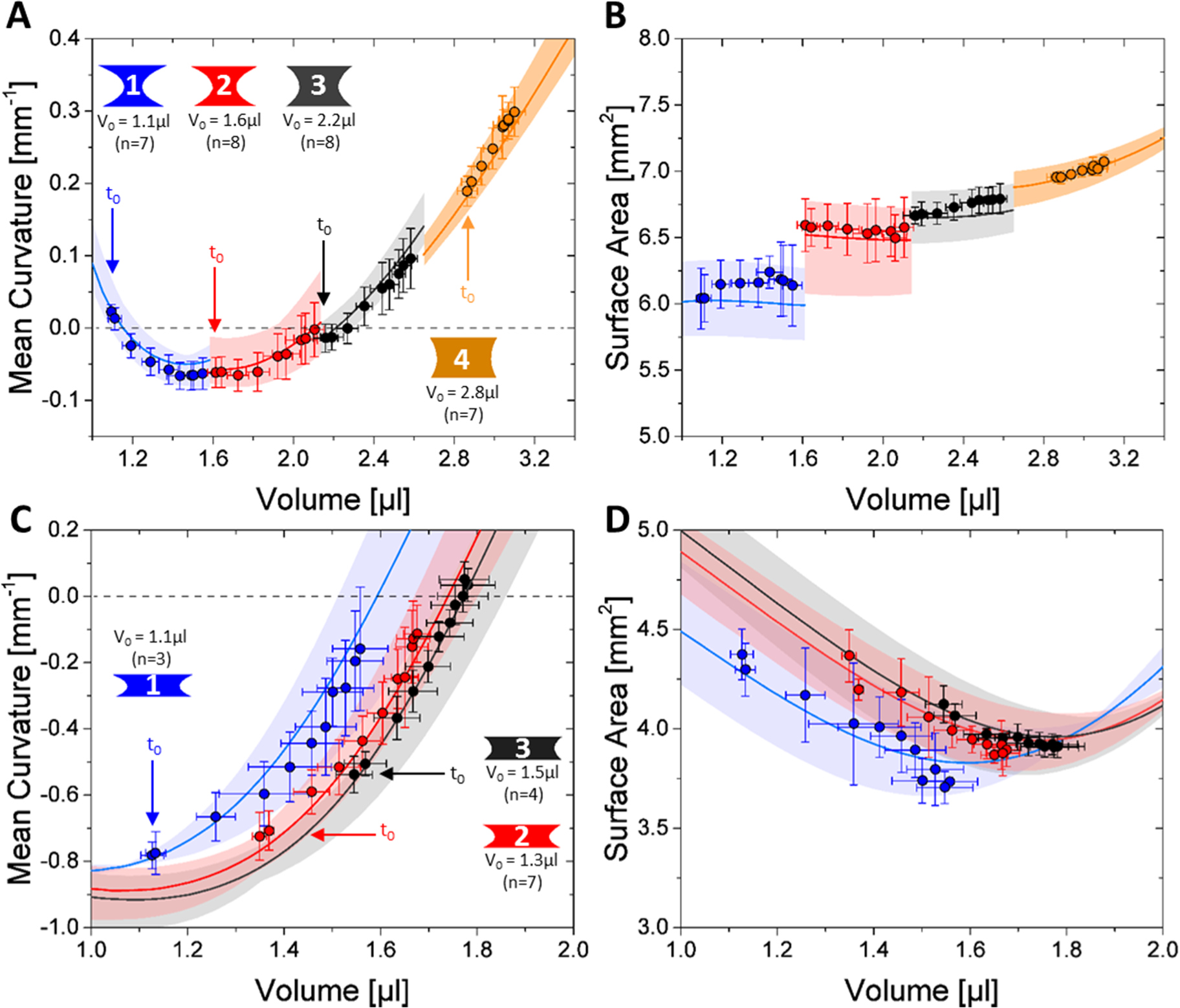
Tissues behave like liquids when constrained by curved surfaces. (A-D) show the evolution of the surface mean curvature (A, C) and surface area (B, D) of the tissue interfaces as a function of total volume (capillary bridge volume V_0_ plus tissue volume) for the two different capillary bridge shapes shown in Figure 1. (A, B) are tissues grown on capillary bridge surfaces with a height of ~1.2mm and top/bottom radius of ~1mm with initial volumes of 1.1μl (Size 1; blue), 1.6μl (Size 2; red), 2.2μl (Size 3; black), and 2.8μl (Size 4; orange) respectively. Colored areas delineate the theoretical predictions of liquid interfaces of same dimension based on the scaffold geometries obtained from experiment. Colored curves are the corresponding mean values. (C, D) are tissues grown on capillary bridge surfaces with a height of ~0.7mm and top/bottom radius of ~1mm with initial volumes of 1.1μl (Size 1; blue), 1.3μl (Size 2; red), and 1.5μl (Size 3; black), respectively. Curves are color-coded according to the corresponding theoretical predictions of the liquid interface. See supplemental material for the full data.

We next investigated whether the kinetics of tissue growth is controlled by surface curvature. By studying various CB shapes, we found that the tissue volume formed after 32 days depends on the CB neck radius, with higher growth rates observed for thin necked CBs (Fig. 3A). In all cases the tissue growth rate showed a similar behavior as a function of time: it gradually increased until reaching a maximum after about two weeks of tissue culture, and then slowed down at later time points. As a consequence, data obtained from different experiments with various CB geometries can be collapsed to one master curve by rescaling the tissue volume V^T^ with the volume at the end of the experiment (V^T^*) (Fig. 3B). To account for differences in initial cell number after seeding onto the scaffold, we further normalized the final tissue volume (V^T^*) by the initial surface area (A_0_) which yields the average final tissue thickness (V^T^*/A_0_). The plot in Fig. 3C reveals that the final tissue thickness correlates with the minimum principle curvature (*k_min_*, concave curvature) of the scaffold, in agreement with previous observations on cylindrical pores [8]. To assess the time dependent slowdown of tissue growth, we normalized the growth rates by the current tissue volume to obtain the exponential growth rate (dV^T^/dt)/V^T^ = d(ln(V^T^/V^T,0^)/dt, plotted in Fig. 3D. Exponential growth would correspond to constant d(ln(V^T^/V^T,0^)/dt and be expected if the rate of cell division was constant over time. This exponential growth rate (or volume strain rate, in mechanical terminology) decreases according to a master curve independent of geometry (Fig. 3D). This suggests that it is related to biochemical signaling or tissue maturation intrinsic to osteoblasts [25] but independent of tissue shape.

**Figure 3:**
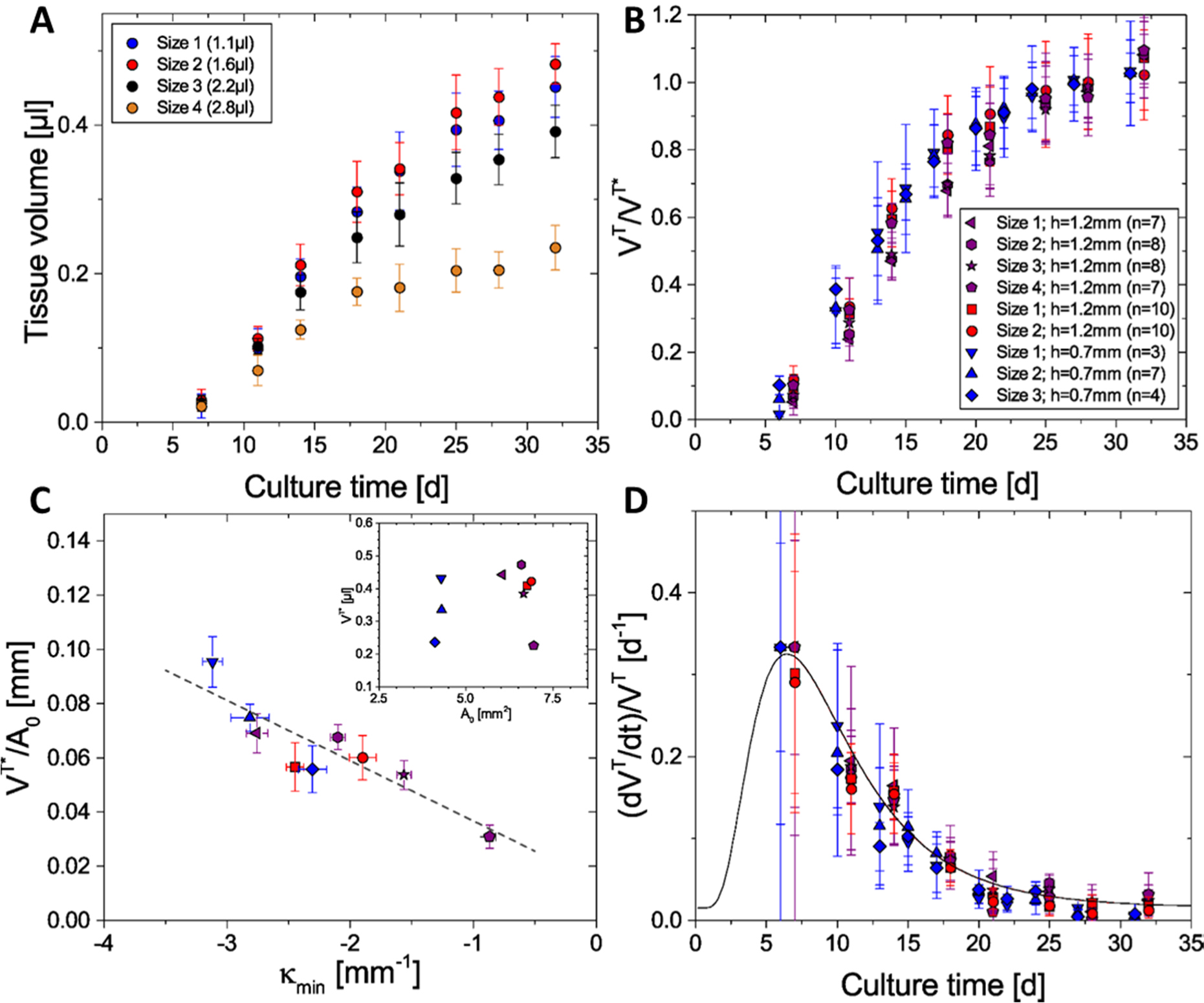
The kinetics of tissue growth depend on the initial curvature. (A) shows the measured tissue volumes for the 4 different capillary bridges presented in Figure 1 (F-I) with initial volumes of 1.1μl (blue, Size 1), 1.6μl (red; Size 2), 2.2μl (black; Size 3), and 2.8μl (yellow; Size 4) respectively. (B) shows a rescaling of the current tissue volume V^T^ with the maximum tissue volume V^T^* (average of day 24 to 32), for 3 independent experiments and different capillary bridge shapes. Data points are color-coded according to the experiment whereby each symbol represents a different capillary bridge shape. (C) shows the average tissue thickness V^T*^/A_0_ as a function of initial minimum principle curvature κ_min_ and a linear regression (grey line, R=-0.936). Inset shows the initial surface area A_0_. (D) shows the rate of growth per tissue volume as a function of culture time, which can be fitted with a decay function following a standard log-normal distribution (black curve).

Finally we investigated the influence of surface curvature on the tissue micro-structure using 3D Light Sheet Florescence Microscopy and actin staining of the tissue. This revealed the formation of chiral actin fiber patterns that exhibit a clear left-handedness (Fig. 4A) in all examined samples (n=12; see suppl. Material). The actin fibers follow locally straight line paths over the curved surface and consistently spiral around the sample at ~60° with respect to the circumferential direction at the neck (Fig. 4B), nearly independently of the capillary bridge shape (Fig. 4C). To interpret this, we investigated three possible hypotheses: First, surface stress in doubly curved surfaces is not homogeneous with respect to the in-plane stress direction. For surfaces with negative Gaussian curvature, the local stress tensor can be decomposed into two normal components where one component has zero stress and the other (perpendicular to it) carries a tangential load. This would mean that the complete surface stress could be transferred just by pulling along this tangential direction with no load perpendicular to it (suppl. Material, membrane-theory). The red line in Fig. 4C corresponds to this favored loading direction. The curve is not defined at small volumes where the mean curvature is such that no zero stress direction exists. Second, we hypothesize that spindle-shaped cells align with the direction of zero curvature (asymptotic directions, suppl. Material). This is inspired from recent observations where fibroblast cells have been shown to behave like active nematic liquid crystals on flat surfaces [26]. This would suggest that cells minimize their free energy by aligning with locally straight directions (Fig. 4C, blue-curve) to obtain an optimal packing on the surface. Third, we approximated the doubly curved surface near the neck by a hyperboloid that is actually a ruled surface generated by a straight line moving through space (suppl. Material). The black line in Fig. 4C corresponds to the direction of the generating straight line. However, none of these three hypotheses gives a perfect fit to the data (Fig. 4C). The red line (Hypothesis 1) is not even defined over the whole range where the chiral patterns were observed. This seems to rule out a purely mechanical origin of the phenomenon and rather suggests liquid-crystal like behavior. This long-range collective cell-alignment also determines the orientation of the collagen fibers that are secreted by the cells which has a direct impact on the mechanical properties and load distribution of the developed/mature tissue [27]. The preference for only one of the asymptotic directions is unresolved but might be related to the intrinsic cell-chirality reported for pre-osteoblasts which exhibit a counter-clockwise orientation when seeded on ring-shaped patterns [28].

**Figure 4:**
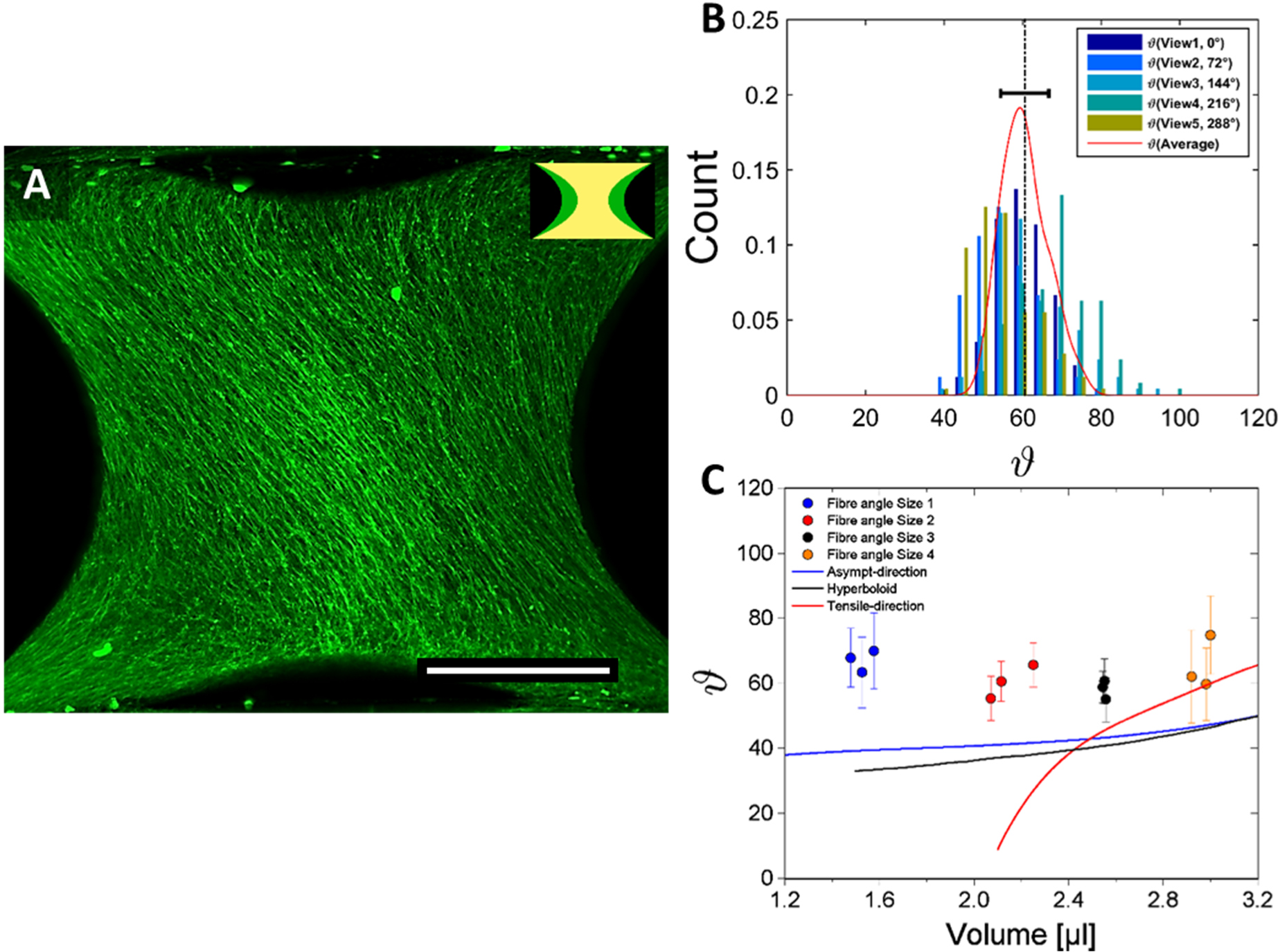
Spontaneous emergence of chiral structures. (A) shows a maximum projections of tissue grown on a capillary bridge with an initial volume of 1.1 μl imaged after 32 days of tissue culture. Tissues were stained for actin and have been visualized using Light-Sheet Fluorescence Microscopy (LSFM). Actin fibres are clearly oriented in a particular direction. Scale bar is 500μm. (B) Shows an example of the fibre angle distributions for 5 different views (72° increments) around the sample obtained using FFT analysis. Red curve delineates average fibre-angle-distribution over all views and dashed line indicates mean value. (C) shows the mean fibre angle distributions as a function of total volume for 4 different sample sizes (color-coded, n=3) compared to asymptotic directions of corresponding capillary bridge surfaces (blue curve) and to hyperboloid surfaces (black curve). Red curve shows theoretical predictions of maximum tensile directions obtained using membrane theory.

Our model system gives quantitative evidence that osteoid-like tissues can make use of the physics of fluids to generate complex three-dimensional patterns, corresponding to equilibrium shapes with constant mean curvature. But we also observed a long-range cell-alignment with a chirality of aligned actin filaments that could preconfigure the orientation of collagen fibrils that form similar patterns in osteons [29, 30]. Such large-scale fiber arrangements also occur in heart tissue where myocardial fibers are organized into a helical muscle band structure [31].

The ability to make scaffolds of defined 3D curvature is highly relevant, since many biological surfaces are doubly-curved [32], with examples including trabecular bone [33] and gyroid structures in butterflies [34]. Our findings about the role of fluidity and surface stresses could potentially also contribute to the understanding of more complex tissue structures such as organoids and may motivate innovative scaffold designs for tissue engineering applications.

